# β-Coronaviruses use lysosomal organelles for cellular egress

**DOI:** 10.1101/2020.07.25.192310

**Authors:** S Ghosh, TA Dellibovi-Ragheb, E Pak, Q Qiu, M Fisher, PM Takvorian, C Bleck, V Hsu, AR Fehr, S Perlman, SR Achar, MR Straus, GR Whittaker, CAM de Haan, G Altan-Bonnet, N Altan-Bonnet

## Abstract

β-Coronaviruses are a family of positive-strand enveloped RNA viruses that include the severe acute respiratory syndrome-CoV2 (SARS-CoV2). While much is known regarding their cellular entry and replication pathways, their mode of egress remains uncertain; however, this is assumed to be via the biosynthetic secretory pathway by analogy to other enveloped viruses. Using imaging methodologies in combination with virus-specific reporters, we demonstrate that β-Coronaviruses utilize lysosomal trafficking for egress from cells. This pathway is regulated by the Arf-like small GTPase Arl8b; thus, virus egress is insensitive to inhibitors of the biosynthetic secretory pathway. Coronavirus infection results in lysosome deacidification, inactivation of lysosomal degradation and disruption of antigen presentation pathways. This coronavirus-induced exploitation of lysosomes provides insights into the cellular and immunological abnormalities observed in patients and suggests new therapeutic modalities.

## Introduction

β-Coronaviruses are positive-strand enveloped RNA viruses that comprise one of the 4 genera of the *Coronaviridae* family of viruses. β-Coronaviruses infect humans and other mammals with infections resulting in a range of diseases with considerable morbidity and mortality. In late 2019, one member, the SARS - CoV2, originating in bats, spread to humans and caused a world-wide ongoing pandemic that up to now has killed over 4 million people (Lu et al., 2020).

The ability of these viruses to infect many different cell types including those of the pulmonary, cardiovascular, hepatic, gastrointestinal, central nervous and immune systems results in complex multi-organ disease manifestations that can vary from individual to individual (Puelles et al., 2020; Ziegler et al., 2020). Especially with regards to the immune system, the virus appears to deregulate the traditional innate and adaptive immune responses to pathogens (Vardhana and Wolchok 2020). Currently there is no cure, the antiviral treatment options are few (Williamson et al., 2020) and it is far from clear if there can even be a lasting immune response generated to infection by natural means or through vaccine administration (Long et al., 2020).

One of the major reasons for lack of anti-viral therapies is the paucity of knowledge regarding the β-Coronavirus-host cell interface. Once the viral envelope fuses with the plasma membrane and/or endosome membranes, and the ∼30kB viral RNA genome is released into the cytosol, it becomes translated into non-structural and structural proteins. The non-structural proteins assemble on ER-derived membranes and replicate the viral RNA (Snijder et al., 2006; Snijder et al., 2020). While some amount of molecular detail is known regarding coronavirus attachment to the cell, receptor binding, envelope fusion and replication, very little is known regarding how the newly assembled coronaviruses exit the cells and spread to others (Fung et al., 2019).

It is largely accepted that the egress pathway for all β-Coronaviruses starts with the newly synthesized viral genomic RNA molecules coated with viral N proteins budding into the lumen of the endoplasmic reticulum (ER) and the ER-Golgi intermediate compartment (ERGIC) (McBride et al., 2007; Cohen et al., 2011; Perrier et al., 2019). This results in the viral genomic materials getting enveloped with host membranes containing the viral M, E and S (i.e. spike) structural transmembrane proteins (de Haan et al., 1998; Siu et al., 2008; Ruch et al., 2012). Once in the ER/ERGIC lumen, the viruses traffic to the Golgi apparatus and Trans-Golgi Network (TGN) for glycosylation and other post-translational modifications (Oostra et al., 2006; Fung et al., 2018). But after the Golgi/TGN, it has been assumed that coronaviruses use vesicles of the biosynthetic secretory pathway to track to the plasma membrane and egress, similar to other enveloped RNA viruses, such as such as hepatitis C, dengue, and West Nile (Ravindran et al., 2016; Robinson et al., 2018).

Here we investigated the egress pathway of β-Coronaviruses and found that rather xthan the biosynthetic secretory pathway, these viruses use an atypical lysosome-based, Arl8-dependent exocytic pathway for release into the extracellular environment. We show that GRP78/BIP an ER-chaperone, that facilitates coronavirus infectivity (Chu et al., 2018; Ha et al., 2020), is co-released with β-coronaviruses through lysosome exocytosis. As a consequence of viral exploitation of lysosomes, we demonstrate that lysosomes are deacidified and lysosomal proteases are inactive. Significantly we show that this perturbation of lysosome physiology impacts functional consequences on the host cell, including disruption of antigen presentation pathways.

## Results

### β-Coronaviruses exit cells independently of the biosynthetic secretory pathway

We began investigating the mechanism of β-Coronavirus egress using mouse hepatitis virus (MHV), as it is the prototype of the family that can be studied under BSL-2 conditions, with intranasal MHV infections in mice inducing pathogenesis similar to SARS, including acute pneumonia, lung injury as well as hepatic and neurological disease (De Albuquerque et al., 2006; Khanolkar et al., 2007; Channappanavar et al., 2016). First, we tested whether MHV used post-Golgi vesicles of the secretory pathway to egress by blocking this pathway with the drug Brefeldin A (BFA) (Lippincott-Schwartz et al., 1989; Miller et al., 1992). We minimized any secondary indirect effects of BFA on viral entry or replication by treating infected cells with BFA between 8- and 14-hour post-inoculation (hr pi) i.e., at the peak virus release **(Figure 1A, black line)**. HeLa-ATCC cells stably expressing murine CEACAM1 (HeLa-mCC1a) were pulsed with virus and at 8hr pi, the culture media was replaced with fresh media with or without BFA (5μgr/ml). After 6hr of incubation (i.e. 14hr pi), both the extracellular media and cell lysates were collected, and their respective viral genome contents were quantified by qPCR. Remarkably, the quantity of virus released in the presence of BFA during egress was almost identical to that in its absence (**Figure 1B, black bars)** and its infectivity when passaged to new cells was unaffected (**Figure S1A**). On the other hand, the secretion of *Gaussia Luciferase*, a transfected reporter that relies on post-Golgi secretory vesicles to be released (Tannous et al., 2009), was completely blocked when BFA was present during infection **(Figure 1B, pink bars)**. These data indicated that the secretory pathway is not essential for viral egress.

**Figure 1.**
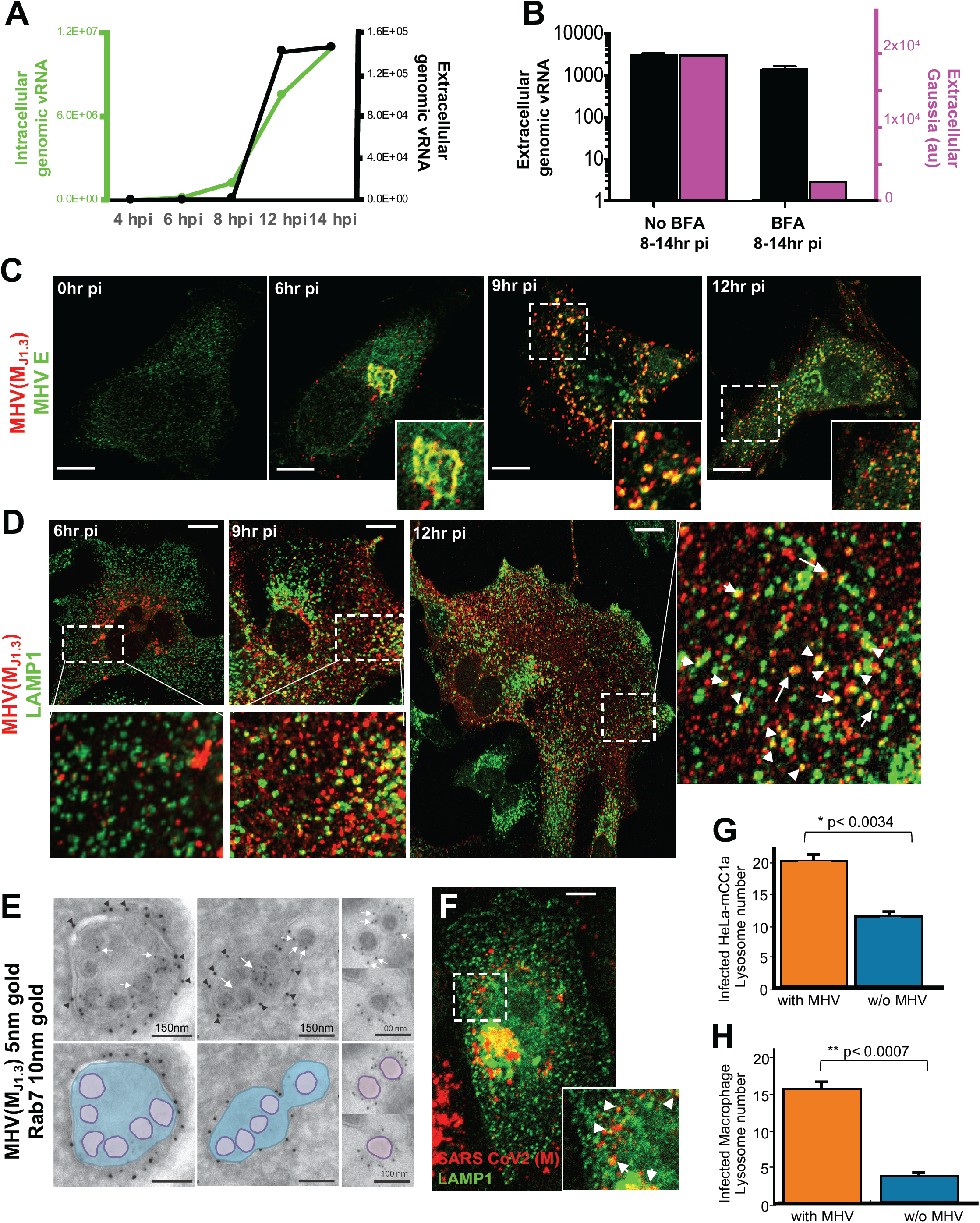
β-Coronaviruses exit cells independently of the biosynthetic secretory pathway. (A) Kinetics of MHV replication and release from HeLa-mCC1a cells. Viral genomic RNA quantified in cell lysates and extracellular medium with QPCR and plotted as fold increase over mock-infection. Experiments done in triplicates. Error bars are SD. (B) Impact of Brefeldin A (5μgr/ml) treatment between 8hr and 14hr pi on MHV release (black bar) or *Gaussia Luciferase* release (pink bar). Extracellular viral genomic RNA was quantified in extracellular medium with QPCR and plotted as fold increase over mock-infection. Experiments done in triplicates. Error bars are SD. (C) HeLa-mCC1a cells infected with MHV, fixed at 0,6,9,12hr pi and coimmunostained with anti-E (green) and anti-MHV (J1.3) (red) antibodies. Scale bar 5μm. (D) HeLa-mCC1a cells infected with MHV, fixed at 6,9,12hr pi and coimmunostained with anti-LAMP1 (green) and anti-MHV (J1.3) (red) antibodies. Arrows point to lysosomes containing M. Scale bar 5μm. (E) Immunoelectron micrographs of Rab7 positive lysosomes (black arrows) containing MHV particles (white arrows). After 12hr MHV infection, HeLa-mCC1a cells expressing Rab7-GFP were co-stained with anti-MHV (J1.3) and anti-GFP primary antibodies followed by 5nm and 10nm-gold coupled secondary antibodies respectively. Virus particles outlined in bottom panel. Scale bars indicated on micrographs. (F) Vero E6 cells infected with SARS-CoV2, fixed at 16hr pi and coimmunostained with anti-LAMP1 (green) and anti-CoV2 M (red) antibodies. Arrows point to lysosomes containing M. Scale bar 2μm. (G) Frequency of LAMP1 positive organelles in MHV-infected HeLa-mCC1a cells (n=16) at 12hr pi. In each cell, lysosomes within a 100μm^2^ region of interest were scored for presence or absence of MHV and plotted. Error bars are SE. (H) Frequency of LAMP1 positive organelles in MHV-infected primary mouse macrophages (n=10) at 12hr pi. In each cell, lysosomes within a 100μm^2^ region of interest were scored for presence or absence of MHV and plotted. Error bars are SE.

We then investigated the spatiotemporal distribution of MHV during infection. The M protein is the most abundant protein in the envelope of β-Coronaviruses and drives virus assembly, membrane curvature and budding into the ER/ERGIC by oligomerizing with itself and with viral RNA, N, E and S proteins (de Haan et al., 2005; Ruch et al., 2012). Immunolabeling cells at peak egress (12hr pi) with the J1.3 monoclonal anti-M antibody (Stohlman et al., 1982; Narayanan et al., 2000), and subsequent immunoelectron microscopy revealed antibody labeling to be concentrated 3-fold more on the envelopes of viral particles than on membranes elsewhere (ER, ERGIC, Golgi etc.) **(Figures S1B and S1C**). While we cannot exclude that J1.3 does not recognize any free M proteins, this indicated that the antibody detected M within the context of assembled particles. In addition, consistent with recognition of assembled virus particles, MHV (M _J1.3_) antibody labeling colocalized with E and S envelope proteins throughout infection (**Figures 1C and S1D**).

We pulsed cells with MHV, fixed at different post-inoculation time intervals, and co-stained with the MHV (M _J1.3_) antibody and antibodies against various host organelle resident proteins. At 6hr pi, consistent with previous reports showing newly synthesized viruses trafficking to the TGN at the early stages of infection (Machamer et al., 2013), MHV (M _J1.3_) labeling was perinuclear **(Figure 1C, 6hr pi)** and colocalized with Golgin 97 and TGN46 by immunofluorescence **(Figures S1E and S1F, 6hr pi)**. Immunogold labeled virus particles could also be observed within ERGIC and Golgi/TGN stacks **(Figure S1G)**.

However, after 9hr pi, the MHV (M _J1.3_) labeling was no longer concentrated at the Golgi/TGN (even though these organelles were still present and intact **[Figures S1E, S1F and S1H]**). Instead it was concentrated in puncta dispersed across the cytoplasm (**Figure 1C, 9 and 12hr pi)**. These puncta appeared to be lysosomes or lysosome-like organelles as they contained lysosomal transmembrane proteins LAMP1 **(Figure 1D, 9 and 12hrpi)**, LAMP2 (not shown), the lysosomal lumenal enzyme cathepsin D **(Figure S1I)** and the lysosomal small GTPase Rab7 (Bucci et al., 2000) **(Figure 1E)**. Significantly, immunoelectron microscopy revealed that the M _J1.3_ labeling reported viral particles in the lysosomes (and not aggregates of M protein or free M protein) **(Figure 1E, white arrows)**. This localization was also not due to reuptake of newly released virus particles as endocytosis was significantly inhibited at this time (**Figures S1J and S1K**). Furthermore, in SARS-CoV2 infected Vero E6 cells, M labeling could also be detected within lysosomes (**Figure 1F**). Quantification of MHV-infected HeLa-mCC1a or primary mouse macrophages revealed that ∼60% and ∼75% respectively of lysosomes contained virus at 12hr pi (**Figures 1G and 1H)**.

### β-Coronaviruses and GRP78/BIP are co-released during infection

We next investigated if any host proteins co-trafficked with MHV to lysosomes during the window of egress. The Golgi resident protein mannosidase II, the TGN resident proteins TGN46 and Golgin 97 **(Figures S1E-S1H**) all remained perinuclear and the mannose-6 phosphate receptor which traffics between TGN and late endosomes (Brown et al., 1986) did not colocalize with MHV in puncta (**Figure S2**). However, the KDEL-Receptor, an ER/Golgi cycling-transmembrane protein that is critical for retrieving escaped ER resident proteins from the Golgi apparatus and the soluble ER luminal chaperone GRP78/BIP (Munro and Pelham 1987) were both found to localize with LAMP1 and MHV during peak virus egress (**Figures 2A-2D**). Moreover, similar to MHV, GRP78/BIP was released into the extracellular media during this time and its release was not due to cell lysis as actin was undetectable in the extracellular medium (**Figure 2E**). Notably BIP release, much like MHV, was insensitive to BFA treatment (**Figures 2F**). Together these data demonstrated that during the time of egress, MHV along with GRP78/BIP, get transferred to lysosomal organelles and released from cells through a route bypassing the BFA sensitive, Golgi/TGN to plasma membrane, biosynthetic secretory pathway.

**Figure 2.**
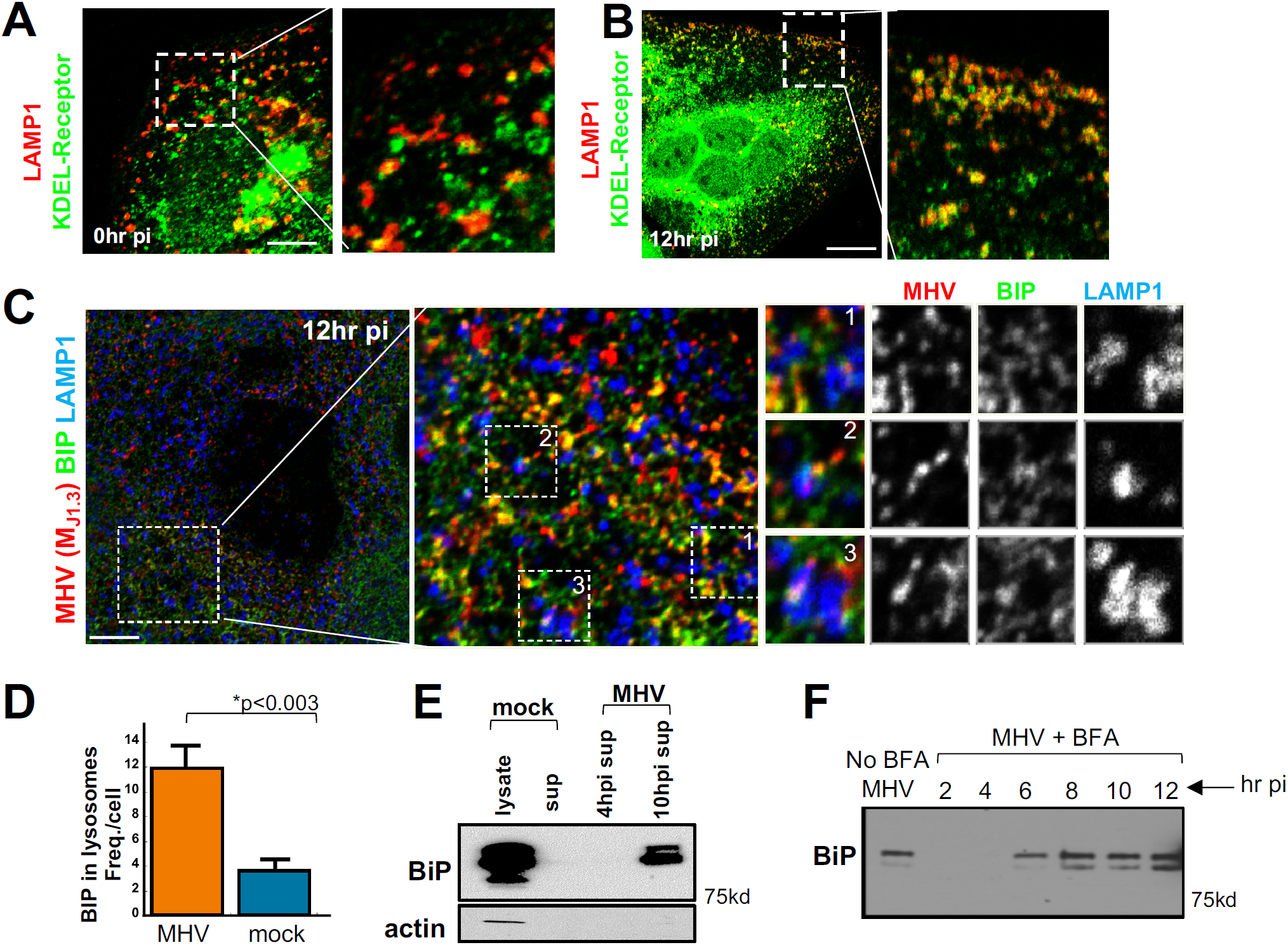
β-Coronaviruses and GRP78/BIP are co-released in infected cells. (A) HeLa-mCC1a cells mock-infected or infected for 12hrs with MHV (B) were fixed and coimmunostained with anti-KDEL-Receptor (green) and anti-MHV (J1.3) (red) antibodies. (A)Scale bar 5μm; (B) Scale bar 10μm. (C) HeLa-mCC1a cells infected with MHV, fixed at 12hr pi and coimmunostained with anti-GRP78/BIP (green), anti-LAMP1 (blue) and anti-MHV (J1.3) (red) antibodies. Scale bar 2μm. (D) Quantification of frequency of GRP78/BIP positive lysosomes in MHV-infected (n=10) and uninfected (n=10) HeLa-mCC1a cells. Error bars are SE. (E) BIP release from MHV infected cells. Cell lysates and extracellular media was collected from mock-infected and MHV-infected cells at 4hr and 14hr pi, processed for SDS-PAGE/Western analysis and probed with anti-GRP78/BIP and actin antibodies. Representative blot from 2 independent experiments is shown. (F) BIP release from MHV infected cells left untreated or treated with BFA(5μg/ml) at 2,4,6,8,10 and 12hr pi. All cell lysates and extracellular media was collected at 14hr pi, processed for SDS-PAGE/Western analysis and probed with anti-GRP78/BIP antibody. Representative blot from 2 independent experiments is shown.

### β-Coronaviruses and GRP78/BIP use an Arl8b dependent lysosome-exocytic pathway for egress

Lysosome exocytosis is a known BFA-insensitive pathway whereby lysosomes traffic to the cell periphery and fuse with the plasma membrane to release their luminal contents (Laulagnier et al., 2011). We conjectured that MHV may be exploiting this route for egress. In support, we found that plasma membrane LAMP1 levels were ∼2.5 fold higher in infected cells **(Figures 3A and 3B)** and cell surface lysosome fusion events (monitored by TIRF imaging of ectopically expressed pHluorin-LAMP1-mCherry) were ∼3-fold enhanced (**Figure 3C)**. Furthermore, immunoelectron micrographs of MHV-infected cells showed numerous Rab7-positive lysosomes containing MHV particles at the plasma membrane with some in the process of fusion **(Figure 3D, black arrows)**.

**Figure 3.**
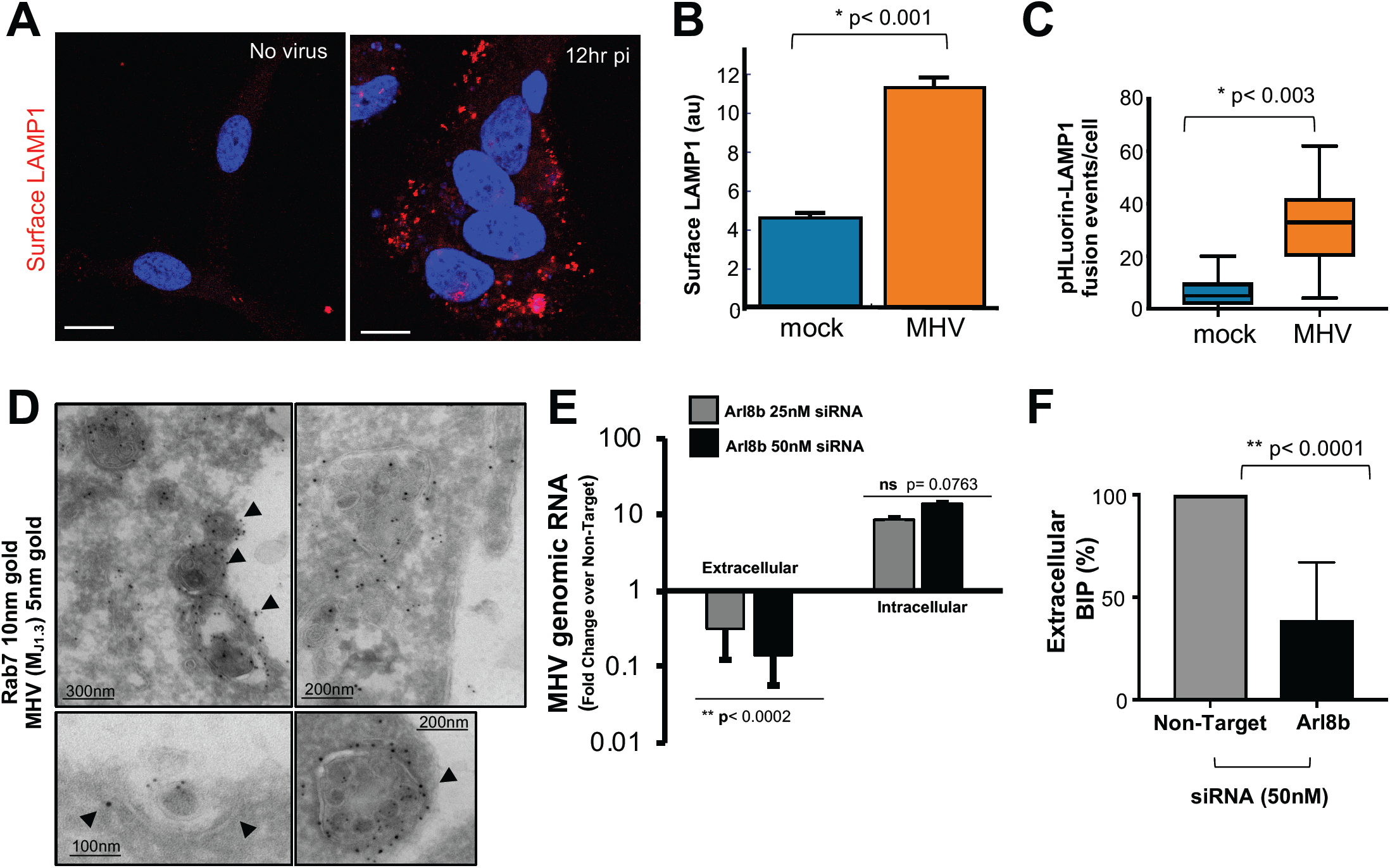
β-Coronaviruses and GRP78/BIP are co-released through Arl8b-dependent lysosome exocytosis. (A) LAMP1 (red) levels on the surface of mock or MHV-infected HeLa-mCC1a cells. Scale bar 10μm. (B) Quantification of LAMP1 staining on the cell surface mock (n=10) or MHV-infected (n=10) HeLa-mCC1a cells. Error bars are SE. (C) Frequency of lysosome plasma membrane fusion events determined with TIRF microscopy on pHLuorin-mCherry expressing mock or MHV-infected (12hr pi) HeLa-mCC1a cells. Minimum 20 cells were imaged for each condition; Error bars are SE. (D) Immunoelectron micrographs of Rab7 positive, MHV-containing lysosomes docked at or fusing with the plasma membrane (arrows). Scale bars are indicated on the micrographs. (E) MHV release in Arl8b siRNA treated HeLa-mCC1a cells. Viral genomes in extracellular media and cell lysates were quantified by qPCR and plotted relative to non-target siRNA treated cells. Representative triplicate data set of MHV genome levels from 4 independent experiments is plotted. Bars are SD. (F) GRP78/BIP release in Arl8b siRNA treated HeLa-mCC1a cells. BIP protein in extracellular media was analyzed by SDS-PAGE/Western, quantified and plotted. Mean of 3 independent experiments is shown. Bars are SE.

We then investigated the role of Arl8b, as a small Ras-like GTPase that localizes exclusively to lysosomes (Xu et al., 2014; Khatter et al., 2015; Michelet et al., 2015; Michelet et al., 2018; Boda et al., 2019) and regulates lysosome movement to the plasma membrane and exocytosis (Michelet et al., 2015). We treated cells with small interfering RNAs (siRNA) targeting Arl8b and obtained ∼50% down regulation of Arl8b in cells **(Figures S3A and S3B)**. Arl8b siRNA treated cells decreased viral release by ∼30-fold compared to non-target siRNA-treated cells and RNA replication was unaffected **(Figure 3E)**. Moreover, Arl8b depletion resulted in >50% decrease in extracellular BIP levels **(Figure 3F)**, confirming that both MHV and GRP78/BIP utilize Arl8b-dependent lysosome exocytic pathways for egress during infection.

### Lysosomes are deacidified and lysosomal enzymes are inactive in β-Coronavirus infected cells

We then assessed the functional consequences of β-coronavirus infection in terms of lysosome function, including SARS-CoV-2 alongside MHV. We used Lysotracker-Red DND99, cell permeable weak base dye that accumulates highly selectively in acidified lysosomes (Sanman et al., 2016). Indeed, labeling cells with the dye prior to fixing with aldehydes and staining with anti-LAMP1 antibodies (without detergents) reveals complete localization of the dye fluorescence to lysosomes (**Figure S4**). HeLa-mCC1a cells and primary mouse macrophages infected with MHV for 12hrs and Vero E6 cells infected with SARS-CoV2 for up to 24hrs were labeled with Lysotracker Red DND99 and aldehyde fixed before imaging. We observed a stark decrease in both the acidity of lysosomes and the number of acidified lysosomes in β-Coronavirus infected cells compared to mock-infected cells (**Figures 4A-4C**). Using the quantifiable live-cell pH dye, Lysosensor Green DND-189, we determined that the mean lysosomal pH in MHV-infected cells was 5.7 with a range between 5.0 to 6.4; whereas in mock-infected cells it was pH 4.7 with range between 4.2 to 5.2 (**Figures 4D)**.

**Figure 4.**
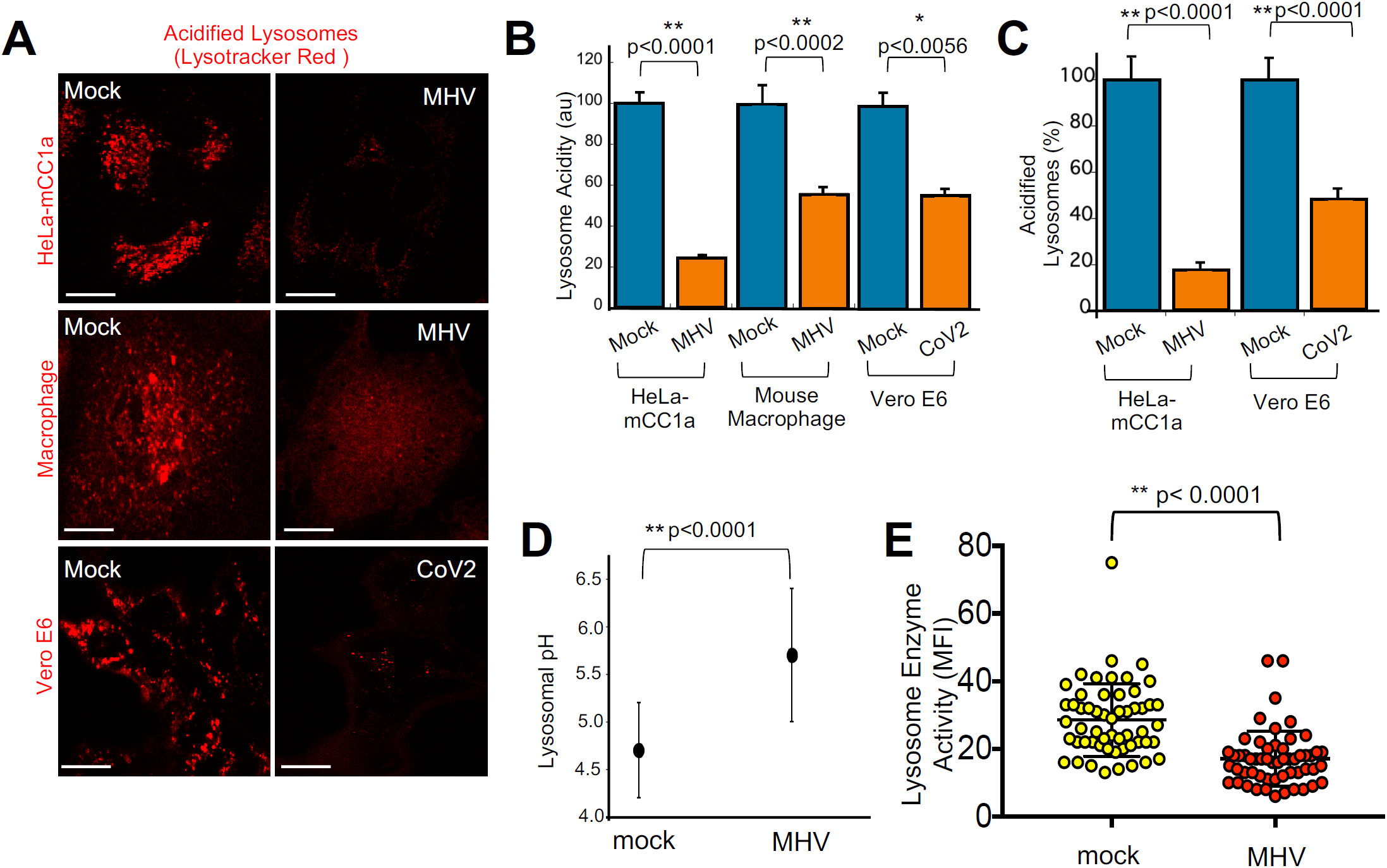
Lysosomes are deacidified and lysosomal enzymes are inactive in β-Coronavirus infected cells. (A) Lysotracker Red DND99 staining of MHV-infected HeLa-mCC1a cells (12hr pi) and primary mouse macrophages (12hr pi); SARS-CoV2-infected Vero E6 cells (16hr pi). Representative images are presented. Scale 5μm for HeLa-CK and macrophage; 10μm for Vero E6 cell. (B) Mean Lysotracker Red fluorescence intensity per lysosome in MHV and CoV2 infected cells (n=20 cells for each condition; 10 lysosomes per cell). Bars are SE. (C) Number of Lysotracker Red positive organelles in mock or CoV2-infected Vero E6 cells (n=30 cells) were quantified and plotted. Bars are SE. (D) Lysosensor Green DND-199 was used to quantify the pH of lysosomes in MHV-infected HeLa-mCC1a cells. Mean fluorescence intensity of 10 lysosomes per mock (n=18) or MHV-infected (n=18) cell was converted to a pH value from calibration of dye. Bars are SD. (E) Lysosome enzyme activity measured in situ in mock- and MHV-infected HeLa-mCC1a cells. Cells were co-incubated with fluorogenic lysosomal enzyme substrate and 10kD dextran-coupled Alexa 555 dye. Plot presents mean fluorescence intensity of the enzyme substrate from similar mean dextran fluorescence intensity lysosomes (n=55 cells each). Bars are SD.

Lysosomal enzymes are optimized for this organelles’ highly acidic pH and even small increases of pH can be sufficient to block protease activity (Sanman et al., 2016; Lie et al., 2019). Given our observation of deacidified lysosomes, we quantified *in situ* lysosomal enzyme activity using self-quenched enzymatic substrates that are taken up by endocytosis, targeted to lysosomes and turn fluorescent upon enzymatic activity (Humphries et al., 2012). As endocytic uptake at peak viral egress was decreased in HeLa-mCC1a cells (**Figures S1J and S1K**), we accounted for this by co-incubating cells with a pH-insensitive fluorophore-coupled dextran, which was endocytosed into lysosomes along with substrate. Mean fluorescence intensity of substrate was quantified in lysosomes with similar mean dextran fluorescence intensity in mock and MHV-infected cells. These measurements revealed that consistent with the observed increased lysosomal pH in MHV-infected cells, lysosome enzyme activities were reduced by ∼40% relative to mock-infected cells **(Figure 4E)**.

### Lysosome-dependent antigen cross-presentation pathways are disrupted in β-Coronavirus infected cells

Finally, we investigated the *functional* consequences of β-coronaviral infection in terms of antigen processing. Myeloid cells rely on active lysosomal degradation of proteins to produce short peptides that are loaded and presented on class I MHC of cells (Trombetta and Mellman 2005). We exposed bone-marrow derived primary macrophages to extracellular chicken Ovalbumin (OVA_1-385_ protein) or to an Ovalbumin-derived class I MHC-restricted oligopeptide (OVA_257-264_ peptide) with or without MHV infection (**Figure 5A and 5B**). First, we measured the endocytosis of fluorescent OVA_1-385_ protein by macrophages and found that it was not significantly affected by MHV infection (**Figure 5C**). Then we measured the amount of OVA antigen being presented by macrophages using the H-2Kb/ OVA_257-264_-responsive OT-1 TCR transgenic T cells (**Figure 5D**). We found that coronaviral infection made macrophages induce stronger T cell activation when presenting OVA_257-264_ peptide, but weaker T cell activation was measured when cross-presenting OVA_1-385_ protein (**Figure 5D**). Such result points out how disrupted lysosomal functions in infected cells alter antigen cross-presentation from protein while possibly boosting presentation from peptide (e.g. by enhancing open conformers of MHC on the surface of cells through increased delivery through lysosome exocytosis)

**Figure 5.**
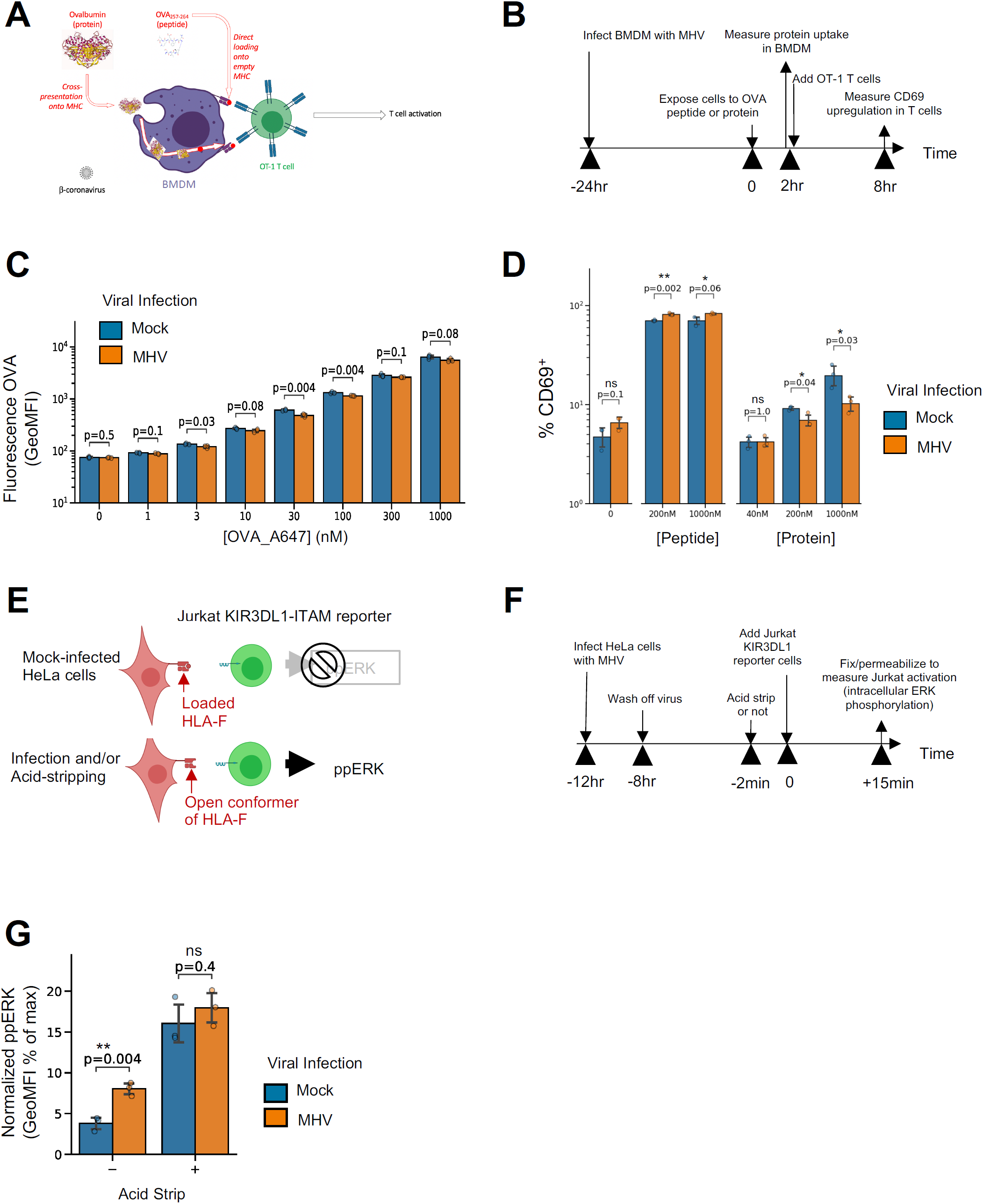
Lysosome-dependent antigen cross-presentation pathways are disrupted in β-Coronavirus infected cells. (A) Sketch and (B) Timeline of assay to test the impact of MHV infection on antigen uptake and cross-presentation. (C) Measurement of fluorescent Ovalbumin uptake by macrophages, with or without prior infection with MHV. (D) Measurement of OVA antigen presentation by macrophages (with or without viral infection) measured as the percentage of CD69+ activated T-cells. (E) Sketch and (F) Timeline of assay to measure the amount of HLA-F open conformers on the surface of infected HeLa-mCC1a cells, using a Jurkat KIR3DL1 reporter cell. (G) ERK phosphorylation in KIR3DL1 reporter cells as a measure of the amount of open HLA-F conformers on the surface of HeLa cells (with or without viral infection). PMA and Null activation are used to normalize ERK phosphorylation, respectively. In all panels in this figure, experiments were done in triplicates. Bars are SE.

We further tested whether coronavirus infection induces the presentation of open conformers of MHC in human cells. Specifically, open conformers of HLA-F could stem from lysosomal dysfunction, serve as activating ligands for NK cells and determine innate immune responses (Goodridge et al., 2013; Garcia-Beltran et al., 2016). We infected HeLa-mCC1a cells with MHV. At 12hr pi we added KIR3DL1-ITAM Jurkat reporter cells (Garcia-Beltran et al., 2016) and measured their ERK phosphorylation after 15min by flow cytometry (**Figure 5E and 5F**). In parallel, we used acid-stripping to open up MHC conformers. We found that MHV infected cells triggered the open-HLA-F-responding Jurkat cells more efficiently, while acid-stripping of cells induced similar (enhanced) response with or without infection (**Figure 5G**). This demonstrated that β-coronaviral infection resulted in enhanced levels of open conformers of HLA-F, another functional immunologically relevant consequence of altered lysosomal activity and cellular stress.

## Discussion

Since the 1960’s, intact coronaviruses have been reported in lysosomes at late stages of infection (Ducatelle and Hoorens 1984) but the significance of these observations remained unknown in terms of viral dynamics. Here we demonstrated that coronaviruses egress from infected cells by tracking a singular path through lysosomes and then to the surface. This is unlike other enveloped RNA viruses whose egress tracks with either the biosynthetic secretory pathway or with direct budding from the plasma membrane.

It remains to be determined if coronavirus trafficking to lysosomes takes place from the Golgi/TGN (a BFA insensitive path (Strous et al., 1993) or from the ER/ERGIC, the latter after retrograde transport back from the Golgi/TGN. Many Golgi and TGN resident proteins, including mannosidase II, Golgin 97, TGN46 and M6PR were absent from the virus-containing lysosomes. In contrast, the KDEL sequence containing chaperone GRP78/BIP, along with the KDEL-receptor were both present. GRP78/BIP facilitates MERS and SARS infectivity (Chu et al., 2018) and has been shown to bind the S protein of SARS-CoV2 (Ha et al., 2020). It was assumed that GRP78/BIP-coronavirus interaction takes place outside the cell (Ha et al., 2020), but our findings here indicate that it is likely taking place already within the host cell during viral egress, since GRP78/BIP is colocalized with coronaviruses in lysosomes and its release to the extracellular environment is Arl8b dependent. Thus, the viruses released maybe primed for best infectivity by being released in complex with GRP78/BIP.

Lysosomal proteolytic enzyme activities are central to many critical cellular processes including autophagy, cell motility, cholesterol metabolism, release of cell killing enzymes by T-cells, pathogen degradation by macrophages and self/non-self antigen presentation by all cells. Lysosome acidification is required for lysosomal enzymatic activities and even a small increase in pH is sufficient to inhibit these enzymes and stop their critical biological functions. We have found that lysosomes are significantly deacidified in coronavirus infected cells and consistent with this measured a significant reduction in lysosomal enzyme activity. The underlying molecular mechanism for the deacidification remains to be investigated but it has been noted that when lysosomes are overwhelmed with cargo - such as they would be in coronavirus infected cells-lysosome deacidification can take place (Ballabio and Bonifacino 2020). In addition, the envelope protein E has been shown to be capable of forming ion channels when ectopically expressed in cells and this may be sufficient to perturb lysosomal proton pumping activities (Ruch and Machamer 2012).

Additionally, our findings indicated that the altered lysosomal function of coronavirus-infected cells resulted in perturbation of antigen presentation and altered immune responses. The flurry of studies triggered by the COVID-19 pandemic has pointed out how unusual and problematic immune responses against coronaviruses can be (Vardhana and Wolchok 2020). Indeed, clinicians as well as basic immunologists have difficulties reconciling observations (e.g. the delayed and erratic macrophage-driven cytokine release syndrome and the severe lymphopenia of CD4+ T and NK lymphocytes) with our current knowledge of immune responses against viruses or cancers (Vardhana and Wolchok 2020). Our findings on the cellular biology of coronavirus and their functional consequence in terms of altered antigen presentation may open new research avenues e.g. focusing on NK cell response against coronavirus-infected cells. The relevance of KIR3DS1-expressing NK cells in delaying the progression to AIDS in HIV-1-infected patients (Martin et al., 2002) or exacerbating the severity of H1N1 infections points (Arando-Romo et al., 2012) out how our findings of increased presentation of open conformers of HLA-F (a known ligand for activating KIR3DS1 and inhibiting KIR3DL1 receptors) could have both positive or negative impact on immune responses against coronavirus infection (Garcia-Beltran et al., 2016).

Finally, coronavirus egress through lysosomal tracking may perturb another innate immune response. Coronaviruses may have evolved to use this egress pathway in order to not only disrupt antigen presentation but also to disrupt endo-lysosomal Toll-like receptor signaling -e.g TLR3 and their binding to double-stranded RNAs, which requires acidification (De Bouteiller et al., 2005). Further studies will need to focus on the impact of such alteration of TLR signaling on inflammatory responses (e.g. cytokine release) triggered by coronavirus infection.

To conclude, our findings open up new avenues to mitigate coronavirus infections, by targeting regulators of lysosomal trafficking such as Arl8b, by reversing deacidification and by enhancing immune cells geared to detect lysosomal malfunction.

## Acknowledgements

NAB and GAB wrote manuscript. NAB, GAB, SG, and TDR designed all experiments. SG, TDR, PT, CB, SA, SG, NAB and GAB performed all experiments. NAB and GAB thank Mary Carrington (NCI) for critical discussion and reagents regarding antigen presentation studies. NAB, SG, TDR, EP, QQ, MF and CB were supported with NHLBI/NIH; GAB and SRA were supported with NCI/NIH intramural funds. PMT was supported by NIH R01 A1091985-05; SP by NIH R01 NS36592 and AF by F32-AI113973; VH by NIH R37GM058615; GW by NIH R01AI35270.

## Supplementary Materials (Supplementary Figure Legends, Materials, Methods)

### Supplementary Figure Legends

**Figure S1(related to Figure 1).**
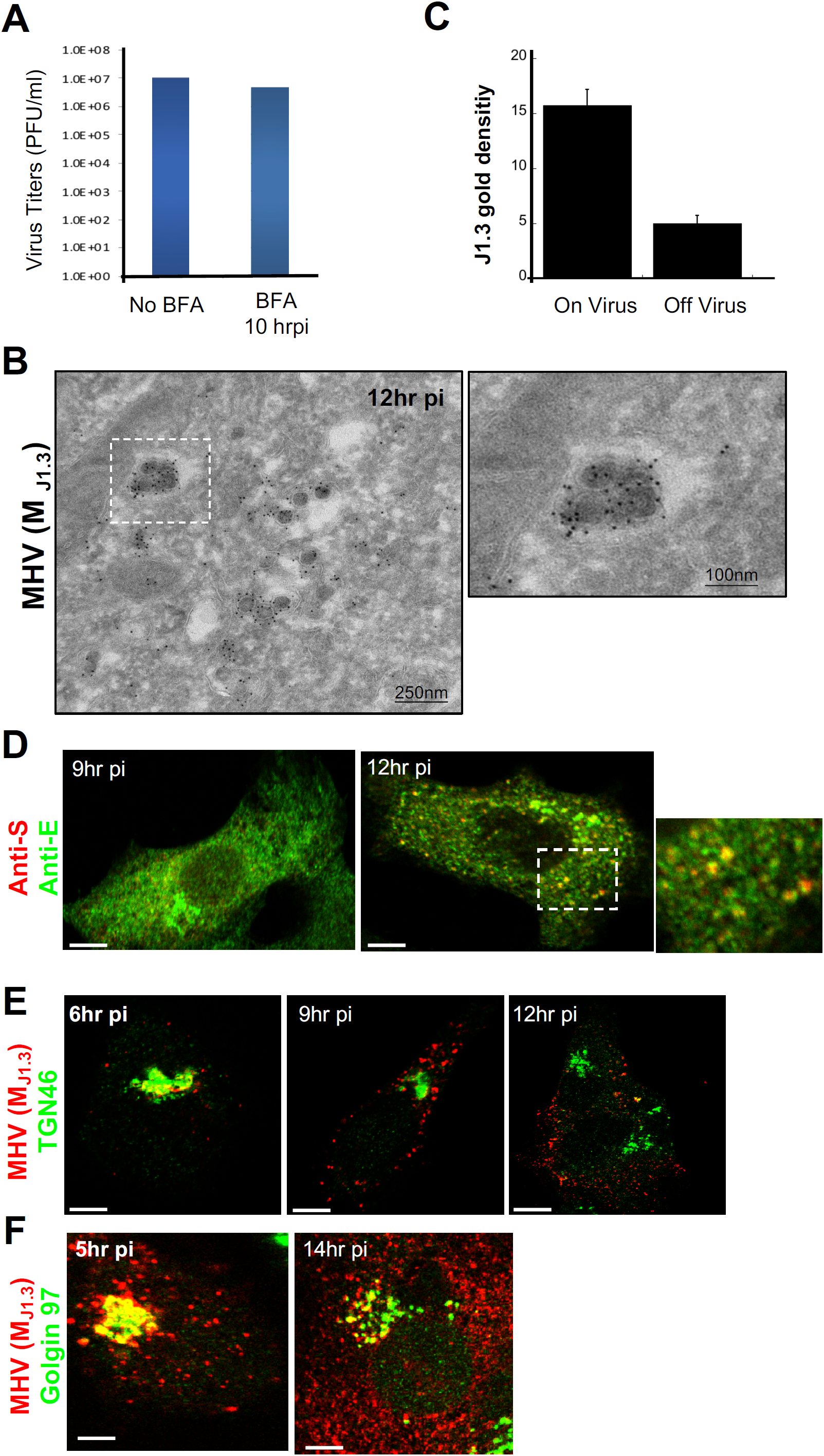
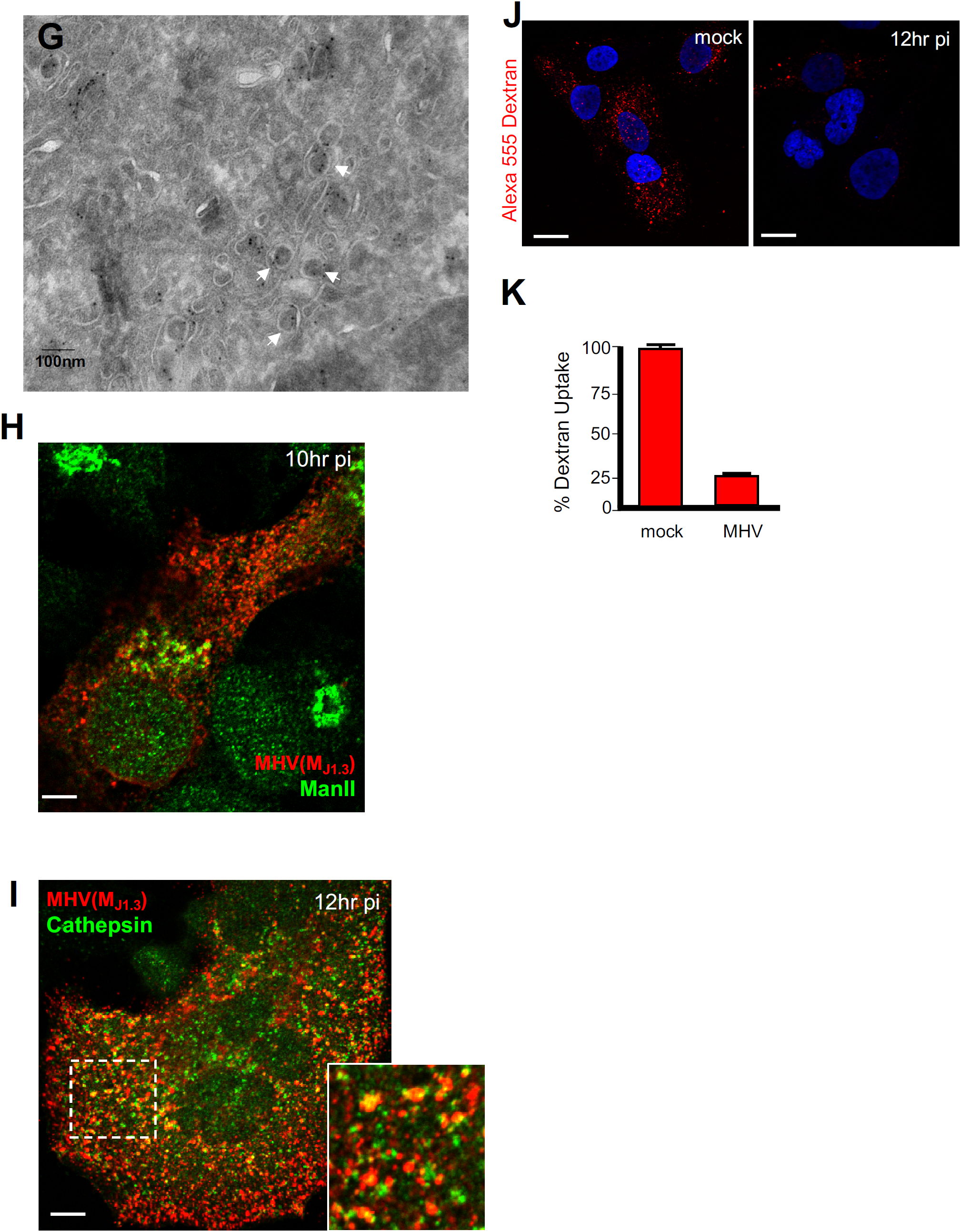
A. Impact of Brefeldin A (5μgr/ml) treatment between 10hr and 14hr pi on MHV release in LR7 mouse cells. Extracellular medium was collected at 14hr pi and used to inoculate new batch of LR7 cells. 24hr post-inoculation PFU/mL was calculated and plotted. B. Immunoelectron micrograph of HeLa-mCC1a cell infected with MHV for 12hr and stained with J1.3 primary antibody and 10nm-gold coupled secondary antibody. The J1.3 labeling is nearly exclusively on the viral particles. C. Quantification of the density of J1.3 anti-M gold over membrane-enveloped virus particles versus cytosolic membranous areas that did not contain discernible virus particles. Gold particles were counted and plotted from regions of interest of equal area in each electron micrograph. Gold particles 5nm. Scale bars indicated on micrographs. D. HeLa-mCC1a cells infected with MHV, fixed at = 9 and 12hr pi and coimmunostained with anti-E(green) and anti-S (red) antibodies. Scale bar 5μm. E. HeLa-mCC1a cells infected with MHV, fixed at 6,9,12hr pi and coimmunostained with anti-TGN46 (green) and anti-MHV (M_J1.3_) (red). Scale bar 5μm. F. HeLa-mCC1a cells infected with MHV, fixed at 5hr and 14hr pi and coimmunostained with anti-Golgin97 (green) and anti-MHV (M_J1.3_) (red). Scale bar 5μm. G. Immunoelectron micrograph of virus particles labeled with anti-MHV (M_J1.3_) (arrows) in Golgi/TGN vesiculo-tubular stacks. Scale bar indicated on micrograph. H. HeLa-mCC1a cells infected with MHV, fixed at 10hr pi and coimmunostained with anti-MHV (M_J1.3_) (red) and anti-mannosidase II (green) antibodies. Scale bar 5μm. I. HeLa-mCC1a cells infected with MHV, fixed at 12hr pi and coimmunostained with anti-Cathepsin (green) and anti-MHV (M_J1.3_) (red) antibodies. Scale bar 5 μm; inset bar 1μm. J. Alexa 555 dextran uptake in mock and MHV infected HeLa-mCC1a cells between 6 and 12hr pi. Representative images collected at 12hr are presented. Scale bars 10μm. K. Quantification of Alexa 555 dextran uptake mock and MHV infected HeLa-mCC1a cells between 6 and 12hr pi.

**Figure S2(related to Figure 2).**
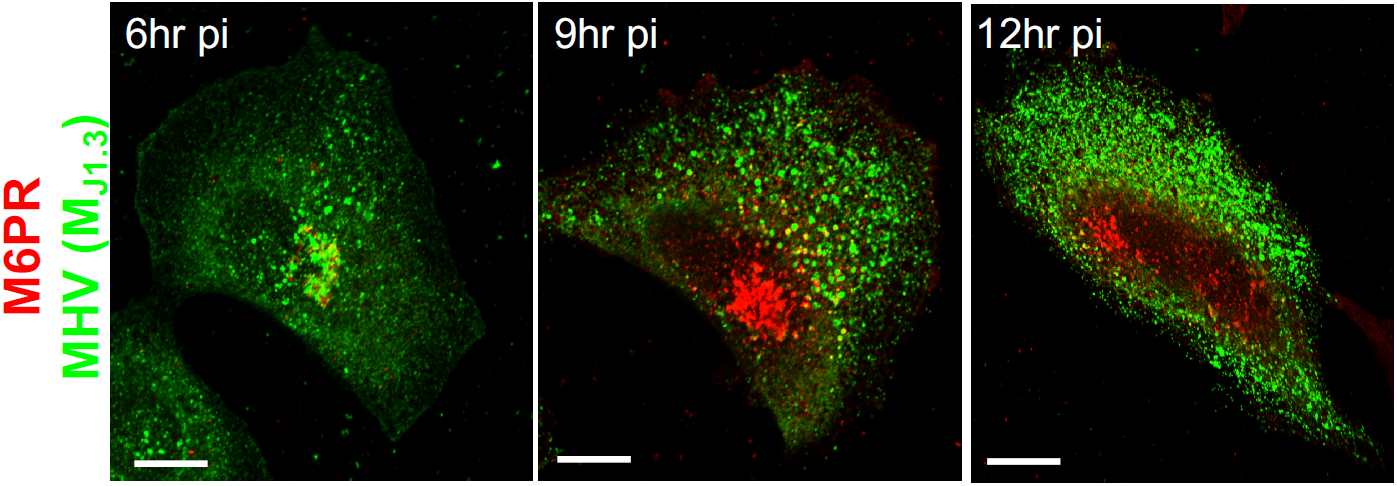
HeLa-mCC1a cells infected with MHV, fixed at 6,9,12hr pi and coimmunostained with anti-MHV (M_J1.3_) (green) and anti-Mannose-6-phosphate receptor (red). Scale bar 5μm.

**Figure S3(related to Figure 3).**
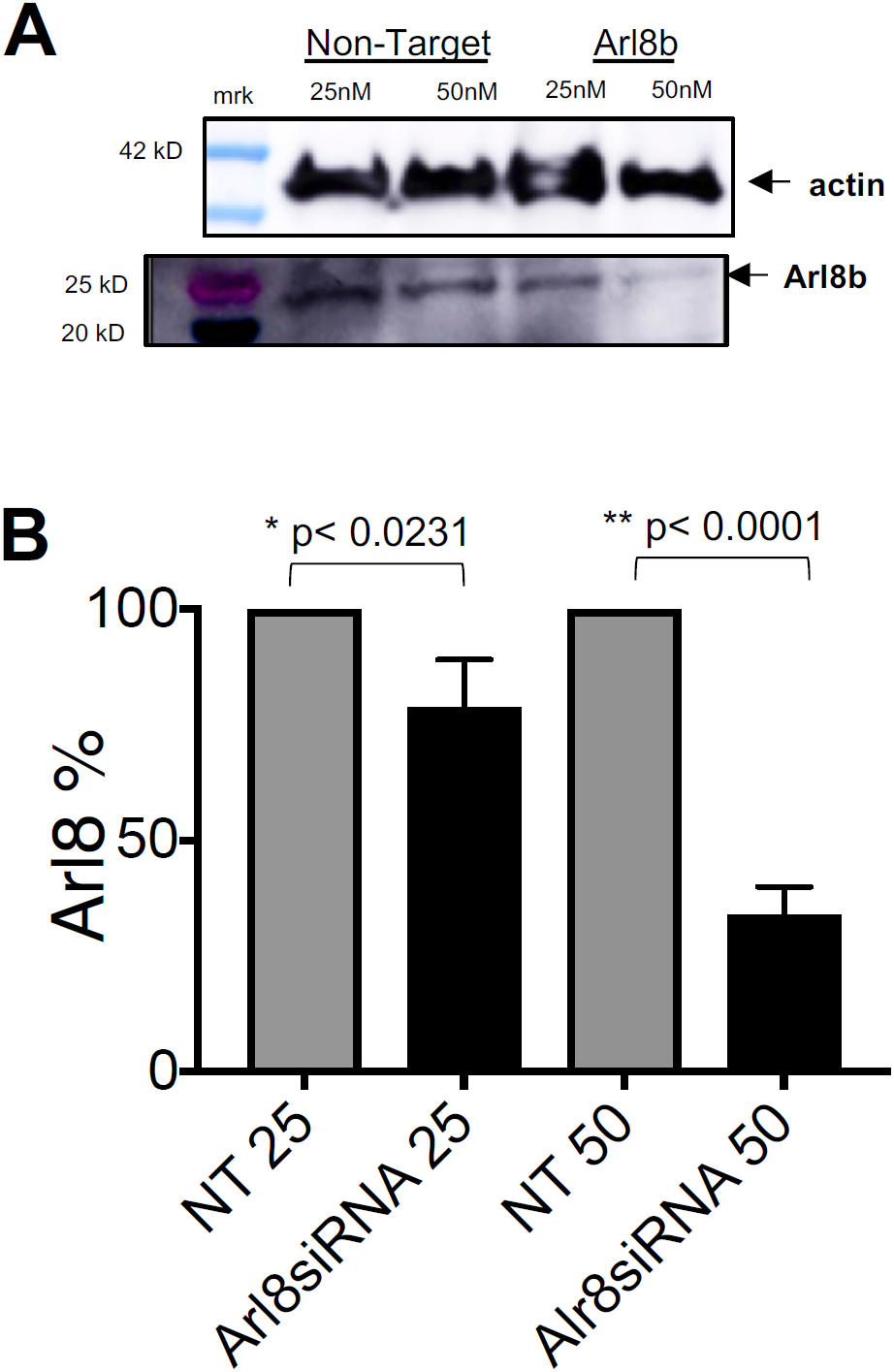
A. Extent of Arl8b depletion after siRNA treatment. HeLa-mCC1a were treated with either 25nM or 50nM of Arl8b or Non-target siRNA for 72hrs. SDS/PAGE Western analysis of cell lysates with anti-Arl8b and anti-actin antibodies was performed. A representative blot is presented. B. Quantification of Arl8b depletion from 4 independent experiments.

**Figure S4(related to Figure 4).**
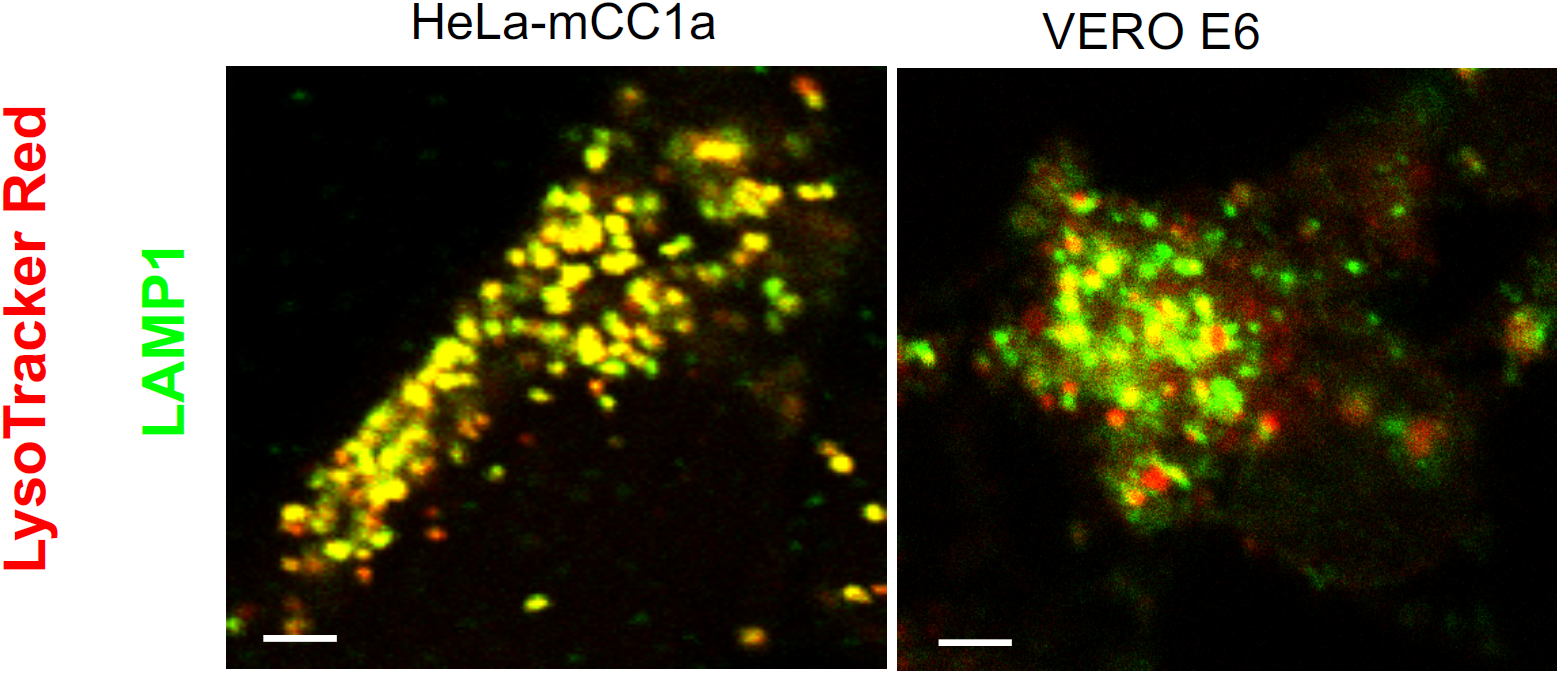
Lysotracker Red-DND99 labels LAMP1 positive lysosomes in HeLa-mCC1a and Vero E6 cells. Scale bar 1μm.

## Supplementary Materials and Methods

### Cells, Viruses, Antibodies, plasmids

KIR3DL1-reporter Jurkat cells were a gift from Dr. Mary Carrington (NCI). pHluorin-LAMP1-mCherry was a kind gift of Harald Stanmark (University of Oslo, Norway). Monoclonal J1.3 antibody was gift from Dr. J Fleming (University of Wisconsin-Madison). Antibodies against TGN46, Golgin 97, GM130, Mannose 6-Phosphate Receptor, cathepsin (Abcam), SARS-CoV2 M (antibodies-online.com) and luminal epitope of LAMP1 (RND Systems) were purchased. Anti-human LAMP1 (H4N3) was purchased from Developmental Studies Hybridoma Bank (University of Iowa). All other antibodies were generated by the authors.

### Virus infections

HeLa-CEACAM1 or LR7 cells were infected with MHV-A59 at MOI 10 for 2-4hrs, washed, and then kept in either complete (with 10%FBS) or serum-free DMEM/high glucose/Pen/Strep for up to 14 hrs.

Vero E6 cells were grown in Millicell EZ 8-well glass slides (Millipore) in infection media (EMEM, 4% FBS (Corning)) to a confluency of 90 – 100%. Cells were then infected with the SARS-CoV2 isolate USA-WA1/2020 at a MOI 1 for 24 hours Mock-infected cells were incubated with infection media only. After 24 hours, supernatants were removed, cells were washed with serum-free media and 100nM or 1µM Lysotracker Red DND99 (Invitrogen) was added to the cells for 1 hour or 10 minutes respectively. Cells were then fixed with 4% paraformaldehyde and washed with DPBS (Corning).

### Immunofluorescence microscopy

Cells were fixed in 3.7% PFA/PBS for 10min; blocked in PBS/10%FBS. All primary and secondary antibody incubations were carried out in PBS/10%FBS supplemented with saponin at 0.2% for 1 hour at room temperature. Cells were rinsed in PBS and mounted with Fluoromount-G (Invitrogen). For surface LAMP1 staining, cells were chilled on ice for 20 minutes and incubated on ice with anti-LAMP1 antibody in PBS for 30 minutes. After rinses with chilled PBS, cells were kept on ice and incubated with secondary antibody in PBS for 30 minutes. Cells were rinsed with chilled PBS, fixed, mounted and imaged.

### Light Microscopy

All microscopy and image acquisition were performed on the LSM780 confocal microscope (Carl Zeiss USA) with a 63X/1.4NA oil objective. Live cells were imaged on a heated stage. Zen software (Carl Zeiss USA) and Image J (NIH) were used for all image analysis.

### Immunoelectron Microscopy

Cells were fixed in 4% formaldehyde and 0.1% glutaraldehyde in 1 x PHEM buffer for 90 min. Cryo-sectioning and immunolabelling were performed as described elsewhere (*1,2*). In brief, ultrathin sections (50–70 nm) from gelatin-embedded and frozen cell pellets were obtained using an FC7/UC7-ultramicrotome (Leica, Vienna, Austria). Immunogold labelling was carried out on thawed sections with anti-GFP (2.5 mg/ml, rabbit, Rockland, 600-401-215) and anti-M (1:50, mouse) antibodies. Mouse primary antibodies were detected with polyclonal rabbit anti-mouse immunoglobulin Gs (0.5 mg/ml). All samples were incubated with 5 or 10 nm protein A gold (1:50, UMC Utrecht University, Utrecht, Netherlands), as described (*1*), and stained/embedded in 4% uranyl acetate/2% methylcellulose mixture (ratio 1:9) (*3,4*).Sections were examined with a JEM-1200EX (JEOL USA) transmission electron microscope (accelerating voltage 80 keV) equipped with a bottom-mounted AMT 6 megapixel digital camera (Advanced Microscopy Techniques Corp).

### RT-PCR Analysis

Cell lysates and supernatants were obtained from specific time points of virus infection and lysed using RNA lysis buffer provided in the RNA isolation kit (Quick-RNA Microprep Kit, Zymo Research, Irvine, CA, Catalog No. R1051). RNA isolation was performed as per the manufacturer’s instructions and cDNA was prepared using Thermo Scientific Maxima First Strand cDNA Synthesis Kit for RT-qPCR (Fisher Scientific, Pittsburgh, PA, Catalog No. FERK1642). RT-PCR was performed using iTaq(tm) Universal SYBR® Green Supermix (BioRad, Hercules, CA, Catalog No. 1725124) in Roche LightCycler 96 system (Roche, Product No. 05815916001). The thermal cycling conditions were composed of a pre-incubation step of 95°C for 90 sec followed by 40 cycles at 95°C for 10s, 54°C for 10s and 72°C for 10s. The samples were run in duplicate for each data point for an experiment. The primers used are mentioned below:

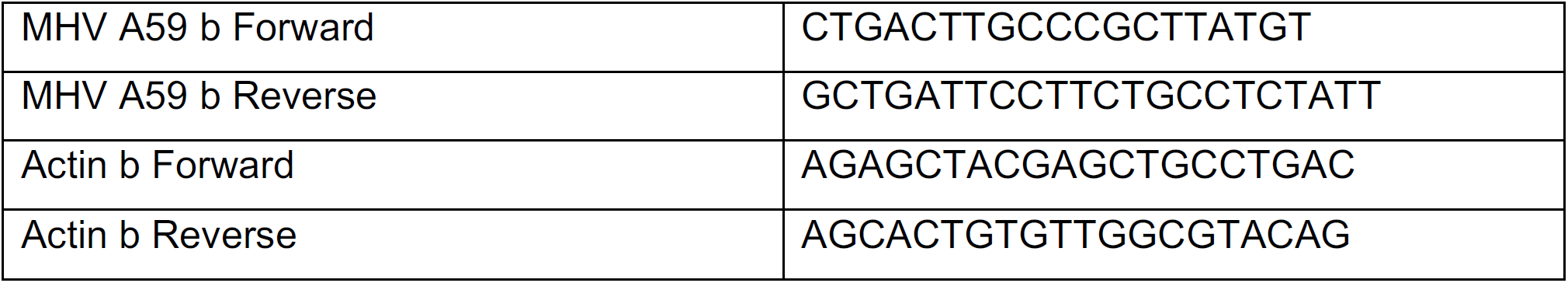

### siRNA treatment

Arl8b and Non-Target siRNA (Horizon Therapeutics) were transfected with Dharmafect 1 (Horizon Therapeutics) and incubated for up to 72 hrs. They were pulsed with MHV for 4hrs, before washing off the virus and switching to serum free media. At 12hr pi, intracellular and extracellular virus was quantified by qPCR. For intracellular RNA levels, cells were lysed with RNA lysis buffer (Zymo Research); for intracellular protein level quantifications cells were scraped and lysed in cell lysis buffer (Invitrogen) containing protease inhibitors. Both intracellular and extracellular proteins were TCA precipitated, acetone washed and suspended in Laemmli gel loading buffer before SDS-PAGE/Western analysis.

### Lysosomal Enzyme Activity

Mock and MHV-infected cells were incubated with Alexa 555 10kD dextran (1mg/ml) and proprietary Lysosome-Specific Self-Quenched Substrate (Abcam Cat No. ab234622) at manufacturers recommended dosage for 1 hour before they were fixed with 4% Paraformaldehyde (PFA) at room temperature for 15 min. Cells were mounted with Fluoromount G (Invitrogen) containing DAPI and imaged with Zeiss LSM780 Confocal Laser Scanning microscope. Images were analyzed using Zen Blue software. Mean fluorescence intensities (MFI) of substrate colocalizing with dextran was measured from each group.

### Lysosensor Green pH measurements

Mock and MHV-infected cells were incubated with Lysosensor Green DND199 (ThermoFisher Scientific) according to manufacturer instructions. Step A: cells were imaged with 488nm laser excitation. Step B: cells were then treated sequentially with potassium buffers of known pH containing 10μM Nigericin; images collected using the same settings as earlier to generate a standard pH curve. Subsequently this standard curve generated in step B was used to convert the fluorescence values collected in step A to pH values.

### Antigen cross-presentation by infected macrophages

Bone-marrow derived macrophages were prepared from femoral aspirates cultured for 7 days in complete RPMI medium with 1nM M-CSF in Fluoroethyl polymer culture bags (Origen). 2.10^2^ cells were harvested and plated on plastic (96-flat-well plate), let to adhere for 2hr, then exposed to MHV (or not) for 24 hr. Macrophages were then pulsed with SIINFEKL peptides or with chicken ovalbumin at varied concentrations for 2 hr, then washed with complete RPMI. C57Bl6 Rag1^-/-^ OT-1 TCR Transgenic mouse splenocytes were then harvested, cleared of their red blood cells by ACK lysis, added and spun onto macrophages (10^2^ cells per well), and incubated for 6 hr. Cell cultures were then harvested using a 15-min trypsin-versene treatment, washed and antibody-stained for flow cytometry (see panel below) with DAPI added just before acquisition. Cells were analyzed using a 5-laser FORTESSA flow cytometer (BD Bioscience) as well as single-stained compensatory UltraComp ebeads (Invitrogen). Data were compensated and processed using FlowJo (TreeStar) and a custom-written Python pipeline (python.org). The level of antigen presentation was quantified using the percentage of live activated T cells (itself estimated as % CD69+ amongst FSC^int^DAPI^-^Vα2^+^ cells).

### Antibody panel for cross-presentation assay

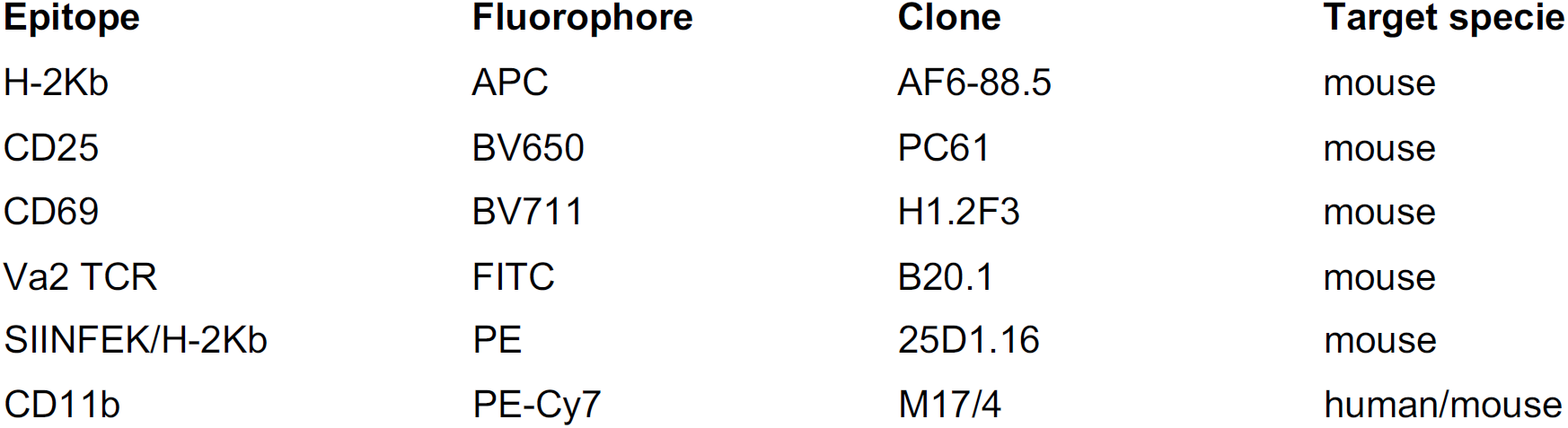

### Endocytosis by infected primary macrophages

Chicken ovalbumin was fluorescently labeled with the Alexa647 dye using a bioconjugation kit (ThermoFisher). Bone-marrow derived macrophages were prepared as described in the previous paragraph. 2.10^2^ cells were harvested, and plated on plastic (96-flat-well plate), let to adhere for 2hr, then exposed to MHV (or not) for 24 hr. Macrophages were then pulsed with varying concentrations of fluorescently-labeled chicken ovalbumin for 2 hr, harvested with a 15-min exposure to a solution of trypsin-versene-EDTA mixture (Lonza), washed with FACS buffer and immediately analyzed by flow cytometry.

### Upregulation of HLA-F open conformers upon MHV infection

HeLa-mCC1a cells were plated and infected (or not) with MHV for 24 hr. Cells were lifted up using a trypsin-versene solution, washed with PBS, incubated for 1min at room temperature with a 0.1M solution of Glycine in PBS (pH adjusted to 2.4) for acid stripping, or with PBS for control, then washed with complete RPMI twice. KIR3DL1-reporter Jurkat cells (*5*) were cultured in complete RPMI and harvested by aspiration. 5.104 HeLa cells were washed with Jurkat cell culture medium then resuspended with 5.104 Jurkat cells, spun at 100g for 15s and incubated at 37°C for 15min (some wells received only Jurkat cells or Jurkat cells and 1µMol of phorbol 12-myristate 13-acetate for negative and positive controls, respectively). Cells were then immediately resuspended with ice-cold 2% PFA for 15 min, permeabilized with ice-cold 90% Methanol for 15 min, washed with FACS buffer and stained for phospho-ERK (E10 clone, Cell Signaling Technology) and anti-mouse secondary antibody (Jackson Immunochemicals). A more detailed protocol can be found in Vogel et al., (*6*). Cells were analyzed using a 5-laser FORTESSA flow cytometer (BD Bioscience). Data were processed using FlowJo (TreeStar) and a custom-written Python pipeline (python.org). The levels of open HLA-F conformers were quantified by monitoring Jurkat cell activation (itself estimated geometric mean of phosphor-ERK staining in FSC^int^ Jurkat cells).

### Statistical Analysis

All graphs were plotted and unpaired two-tailed Student-t Test was performed using the GraphPad Prism 8 software or the SciPy Statistics library in Python. *p* values were considered significant for *p*< 0.05 unless otherwise indicated and denoted as */** and the corresponding values are mentioned in the figures.

## Notes

### Competing Interest Statement

The authors have declared no competing interest.

